# Missing cell types in single-cell references impact deconvolution of bulk data but are detectable

**DOI:** 10.1101/2024.04.25.590992

**Authors:** Adriana Ivich, Natalie R. Davidson, Laurie Grieshober, Weishan Li, Stephanie C. Hicks, Jennifer A. Doherty, Casey S. Greene

**Author notes:** **Corresponding Author:** Casey S. Greene.

## Abstract

Advancements in RNA-sequencing have dramatically expanded our ability to study gene expression profiles of biological samples in bulk tissue and single cells. Deconvolution of bulk data with single-cell references provides the ability to study relative cell-type proportions, but most methods assume a reference is present for every cell type in bulk data. This is not true in all circumstances--cell types can be missing in single-cell profiles for many reasons. In this study, we examine the impact of missing cell types on deconvolution methods. Our experimental designs are simulation-based, using paired single-cell and single-nucleus data, since single-nucleus RNA-sequencing is able to preserve the nucleus of cell types that would otherwise be missing in a single-cell counterpart. These datasets allow us to examine the missing-cell-type phenomenon in deconvolution with realistic proportions. We apply three deconvolution methods that vary from straightforward to state-of-the-art: non-negative least squares, BayesPrism, and CIBERSORTx. We find that the performance of deconvolution methods is influenced by both the number and the similarity of missing cell types, consistent with prior results. Additionally, we find that missing cell-type profiles can be recovered from residuals using a simple non-negative matrix factorization strategy. We expect our simulation strategies and results to provide a starting point for those developing new deconvolution methods and help improve their to better account for the presence of missing cell types. Building off of our findings on simulated data, we then analyzed data from high-grade serous ovarian cancer; a tumor that has regions of highly variable levels of adipocytes dependent on the region from which it is sampled. We observe results consistent with simulation, namely that expression patterns from cell types likely to be missing appear present in residuals. Our results suggests that deconvolution methods should consider the possibility of missing cell types and provide a starting point to address this. Our source code for data simulation and analysis is freely available at https://github.com/greenelab/pred_missing_celltypes.

## Introduction

Gene expression analysis, particularly through RNA-sequencing (RNA-seq), provides a means to characterize transcript levels in complex tissues. Bulk RNA-seq has been widely applied for capturing aggregate snapshots of gene expression within tissues [1, 2]. The advent of single-cell level technologies has brought in a new era, revealing cellular heterogeneity at a higher resolution that was not observable by bulk approaches [3-5].

Previous studies have shown differences in both gene expression and cell composition between single-cell, single-nucleus, and bulk RNA-seq data [4, 6-10]. In some cases, cell dissociation and processing in single-cell RNA-seq could introduce complete or partial cell loss in cell types that are adherent, sensitive, too large, or have low capture efficiency [6-8]. We expected the impact of this loss might be particularly pronounced when attempting to link observations from single-cell and bulk datasets, since certain cell types may be absent from single-cell profiling.

While not directly observable, cellular heterogeneity could be inferred from bulk sample by bulk deconvolution. Deconvolution methods play a pivotal role in unraveling the composition of cell types within the vast amounts of archival and relatively accessible bulk RNA-seq datasets. Estimating cell-type proportions in bulk datasets is key in identifying cell proportion differences and performing accurate differential gene expression analysis at the cell-type level. These deconvolution methods (when reference-based) rely on a single-cell expression reference (referred to as cell reference) to estimate cell-type proportions in bulk [11, 12]. When single-cell RNA-seq experiments are used as ground truth, the reliability of deconvolution is contingent on the completeness of the reference, posing a critical challenge when certain cell types are absent [10].

High-grade serous ovarian cancer (HGSOC) is a fitting instance of the importance of considering absent cell populations when analyzing single-cell versus bulk RNA sequencing data. In HGSOC, key questions exist as to the nature of previously described transcriptomic subtypes [13, 14], and previous studies have raised the possibility that subtypes may be driven by cell-type proportion differences [15, 16]. Additionally, previous work from our group has found an impact of dissociation on adipocyte-related gene sets, revealing lower expression in dissociated bulk RNA-seq samples compared to their undissociated counterparts, suggesting loss of adipocytes during deconvolution [17]. This discrepancy in expression prompts critical questions about the fidelity of deconvolution methods in the face of missing adipocyte information, particularly when such information could hold clinical relevance. The metastatic potential of HGSOC to the omentum [18, 19], which is rich in adipose tissue, underscores the need to discern the implications of missing adipocyte information in deconvolution analyses. Omentally derived samples may contain more adipocytes than samples from other intraperitoneal locations simply because they are derived from a tissue with more adiposity. We hypothesize this would cause omental bulk samples to have higher expression of adipocytes, while the single-cell RNA-seq counterpart would not have these cells. While our work raises this possibility in HGSOC, missing cell types would impact any tissues where certain cell types are difficult to isolate in a single-cell suspension.

Previous studies have explored effects on deconvolution performance of the remaining cell types after removing one cell type at a time [20, 21] from the cell reference in the context of testing the robustness of deconvolution methods [11, 22, 23]. These studies focused on the effect of a cell type being removed from the cell reference on the redistribution of predicted cell proportion. Furthermore, they also found that similarity in gene expression, or correlation of expression, between the cell types missing from the cell reference and the remaining cell types in the cell reference can affect the predicted proportions [22]. These previous studies have only examined the effect of removing one cell type at a time and have only removed cell types that are typically not missing from single-cell RNA-seq studies. Also, previous studies have not attempted to recover the missing cell-type information.

Building upon this foundation, we aim to examine the effect on deconvolution performance when multiple cell types are missing from a reference single-cell dataset used in bulk deconvolution, and whether the missing cell-type information is recoverable. We used a curated set of immune cell types, the PBMC3k dataset provided by 10x Genomics [24], and increased the simulation’s physiological relevance by including white adipose tissue and single-cell RNA-seq data with real missing cell types. Simulated bulk data (referred to as pseudobulks) were generated from these single-cell (and single-nucleus) datasets for deconvolution, incorporating random and realistic proportions with and without Gaussian noise (see Methods for details). We used three deconvolution methods, non-negative least squares (NNLS [25]), CIBERSORTx [12], and BayesPrism [11] in our experiments, and calculated residuals matrices by subtracting recreated bulks from original pseudobulks. We examined whether evidence of the missing cell types was present in residuals by applying non-negative matrix factorization (NMF). We also examined a dataset of matched HGSOC bulk and single-cell RNA-seq [17], in which we expect the single-cell data to lack a cell type expected in the bulks (adipocytes). Through these approaches, we carefully examine the consequences of missing cell types on deconvolution outcomes and evaluate the extent to which it is possible to recover absent cell-type information. We do not present a novel method. Instead, our findings suggest paths to improve deconvolution methods, either by adding an iterative step that uses residuals to identify structure that may align with cell types and model it or by using study-specific references alongside large atlas-based panels.

## Results

### Missing cell-type information is present after deconvolution

We sought to examine whether methods could recover missing cell-type proportions by first using pseudobulks with 5 distinct immune cell types (Figure 1A; see Methods for details). These cell types have variable similarities in their expression profiles, shown in Supplemental Figure 1A. We deconvolved pseudobulks with NNLS, the baseline method, with 0, 1, 2 and 3 missing cell types in the cell reference. When no cell types were missing, NNLS demonstrated strong performance, accurately recovering the real proportions of each cell type (Figure 2A). However, as the number of missing cell types increased, the performance of NNLS gradually declined (Figure 2B-D). To determine whether the missing cell-type proportions were recoverable, we calculated residuals as shown in Figure 1B (See Methods: Residual Calculation). When plotting the factorized residuals against the true proportion for each of cell types present in the pseudobulks, NMF factors were found to be correlated with the proportions of each missing cell type (Figure 2 E, F and G for 1, 2 and 3 missing cell types respectively). The missing cell-type proportions show high Pearson’s correlation and low RMSE values for each of the missing cell types (Fig. 2E-G).

**Figure 1.**
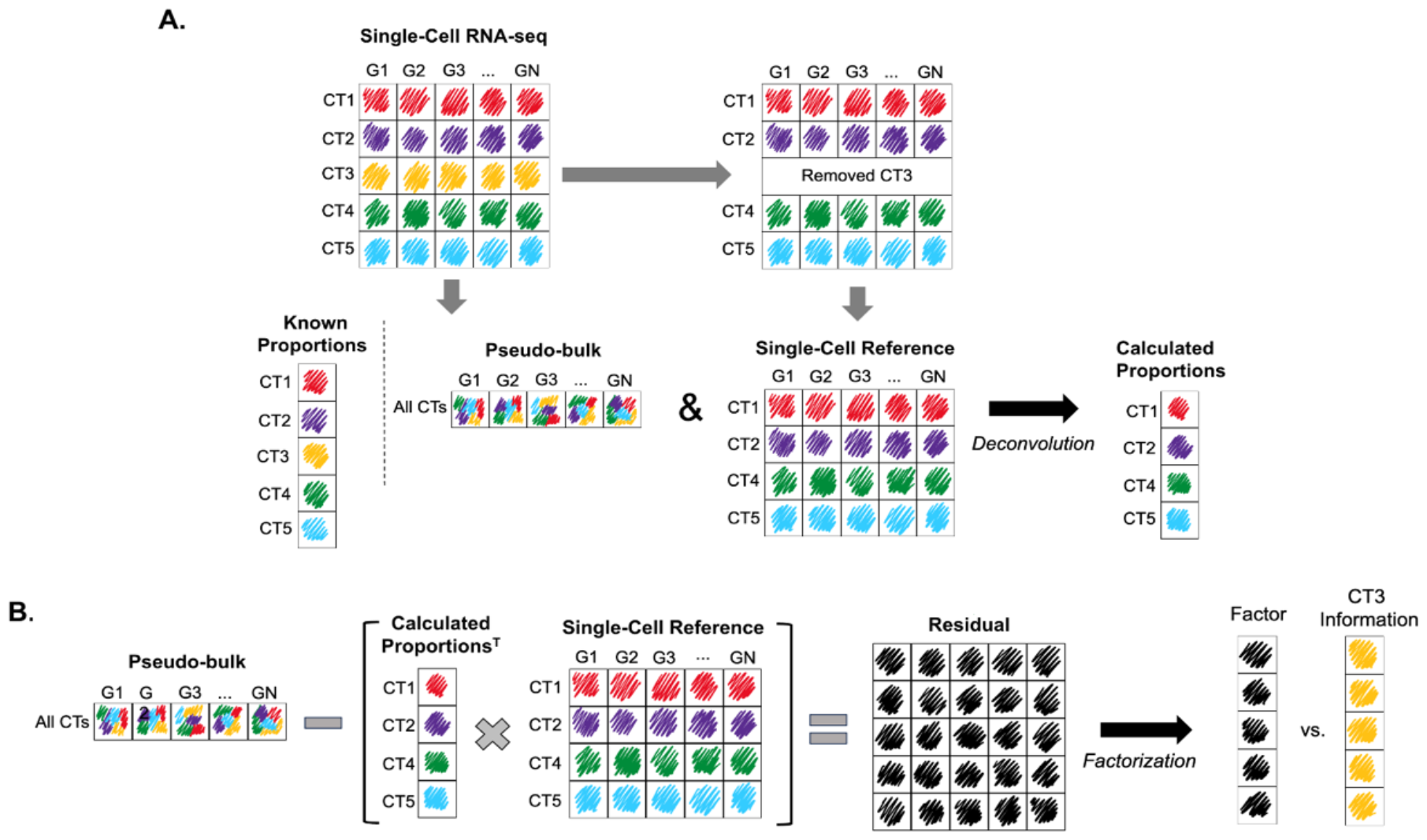
Schematic illustration of experimental design. **A**. Single-cell RNA-seq dataset is used to create pseudobulks (bulks with known proportions). Then, a cell type in the single-cell reference is removed. The pseudobulks are deconvolved with that reference, to obtain the calculated proportions. **B**. The calculated proportions are multiplied by the single-cell reference used, to create a recreated expression matrix. Then, this is subtracted from the original pseudobulks to get the residual matrix, which is factorized to find the missing cell-type’s proportions.

**Figure 2.**
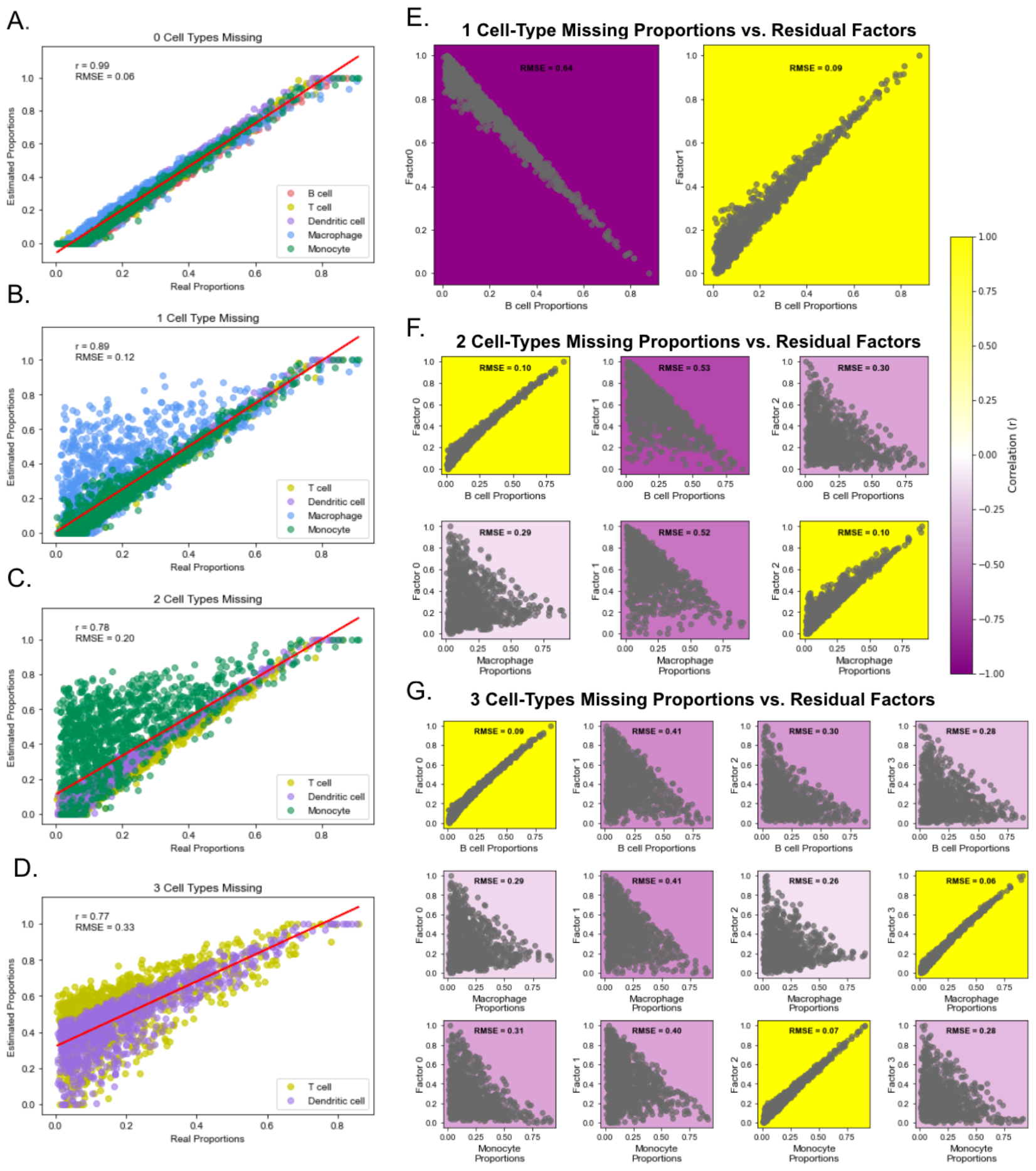
Residual NMF of distinct immune cell types from Single-Nucleus RNA-seq Adipose Tissue using NNLS. The panels on the left (A-D) show the deconvolution performance of NNLS in random-proportioned pseudobulks with **A**. zero cell types missing, **B**. one cell-type missing, **C**. two cell types missing, and **D**. three cell types missing from the NNLS deconvolution cell reference. Pearson’s correlation and RMSE are noted in each plot between the real and calculated proportions. Panels on the right show each missing cell-type’s proportions correlated with each residual’s NMF factor across the number of missing cell type; **E**. reference with one cell type missing, **F**. two cell types missing, and **G**. three cell types missing. Each row represents one of the missing cell types, and each column represents each normalized NMF factor for each panel. The color represents Pearson’s correlation (see color bar), and RMSE is noted in each box.

### Missing cell-type profiles are present in residuals from widely used methods

NNLS is a straightforward method for this problem, but multiple more sophisticated methods are more commonly deployed in practice. We designed a similar experiment to test two more widely used methods, BayesPrism and CIBERSORTx (which is marker gene-based), alongside NNLS. Additionally, we added more biological complexity to our methodology by creating a new set of pseudobulks from the PBMC3k dataset, which contains cell types with more similar expression that our previous section (see Methods for details). This experiment represents a more challenging scenario for deconvolution algorithms.

Specifically, we first removed cell types from the deconvolution cell reference for which the reference profiles were relatively uncorrelated (Supplemental Figure 1B, removed cell types are B cells, FCGR3A Monocytes, and NK T cell, and CD4 T, cumulatively). We analyzed these data as in our first experiment, using BayesPrism and CIBERSORTx in addition to NNLS. For BayesPrism and CIBERSORTx, we used each model’s directives according to the original publication while creating the references [11, 12]. For NNLS, the cell reference is created as in the previous experiment (see Methods for details). We observed similar effects on the performance of deconvolution methods as we increased the number of missing cell-types. Figure 3 shows deconvolution performance for NNLS, BayesPrism, and CIBERSORTx in Panels A-C, respectively.

**Figure 3.**
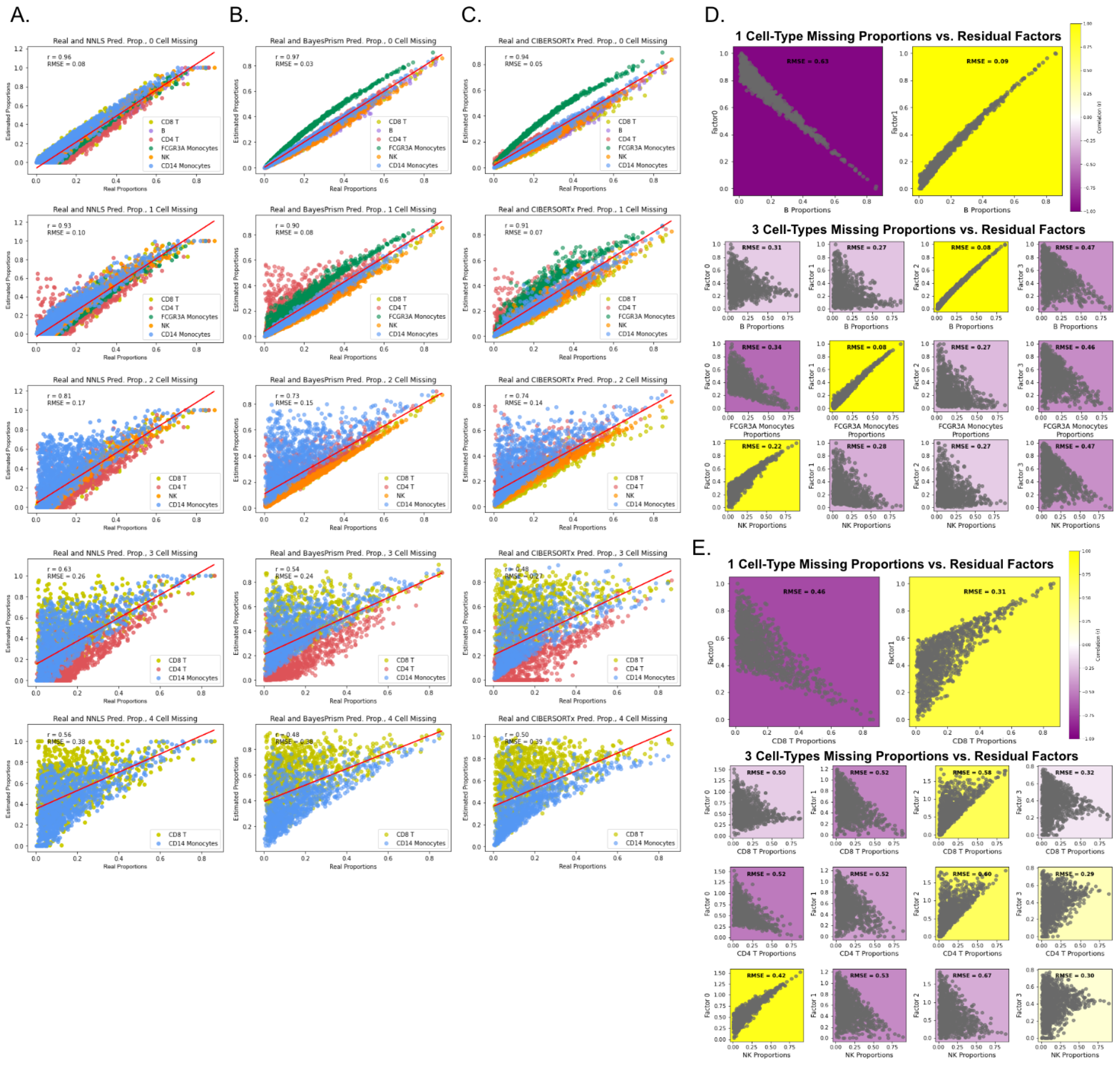
Deconvolution performance across deconvolution methods as we increase the number of missing cell types from the reference (random proportions), and the accuracy of recovering the proportions is dependent on how similar the missing cell types are to the other missing cells in random-proportioned pseudobulks (PBMC3k) deconvolved with NNLS, with residual factorization through NMF. **A**. NNLS, **B**. BayesPrism and **C**. CIBERSORTx performance (real vs. estimated) proportions. First row corresponds to the control (zero missing cell types), second row corresponds to one missing cell type (B cells), then two missing cell types (B cells, FCGR3A Monocytes), three (B cells, FCGR3A Monocytes, NK) up to four missing cell types (B cells, FCGR3A Monocytes, NK, CD4 T cells) in the last row. Each cell’s proportions are colored a different color (legend), and each plot has Pearson’s correlation and RMSE value noted. The panels on the right (D and E) show the correlation between the residual’s factor and each missing cell type, comparing the deletion of **D**. cell types with distinct expression are deleted from the single-cell reference, and the residual is factorized and **E**. cell types with similar expression are deleted from the single-cell reference, and the residual is factorized.

We next determined whether the residuals from these additional methods similarly contained apparent signal from missing cell types. To do this, we calculated and factorized the residuals. We often found factors highly correlated to each of the missing cell types. We observed this pattern irrespective of method (Supplemental Figures 2A, 3A and 4A show NNLS, BayesPrism, and CIBERSORTx, respectively). This suggested that information present in the residuals was likely to represent cell types, raising the possibility that future methods might be able to tap into this latent structure.

We hypothesized that the recoverability of this information would depend on the similarity of the missing cell-type’s expression profiles, both with other removed cell types and cell types in the pseudobulks, as see in previous work [22]. We designed an experiment to examine whether similarity influenced recoverability by removing CD8 T cell, CD4 T cell, NK T cell and FCGR3A Monocytes cells, cumulatively, from the same data.

These cells have relatively more correlated expression with other cell types (Supplemental Figure 1B) compared to the previous removed cells. We found differences between these conditions (Figure 3 D vs. E). Although deconvolution performance was comparable for all deconvolution methods regardless of whether the missing cell types were similar or not (Supplemental Figure 5: A vs. B, and Figure 3 A, B, C vs. Supplemental Figure 6), the recoverability of missing cell-types’ proportions appeared substantially more dependent on the identity of the missing cell type. When we remove very distinct cell types, we observe that the true proportion of the cell type uniquely matches to one NMF factor (Figure 3D). This contrasts with the NMF factors of residuals after more similar cell types are omitted (Figure 3E).

### Matched single-cell and single-nucleus data enable testing with realistic proportions

Our experiments thus far have constructed scenarios in which the missing cell type is removed without regard to what is physiologically likely or realistic. To examine the effect in a more realistic scenario, we sought to identify a condition where we could create profiles that mimic actual bulk/single-cell discrepancies while still retaining a gold standard for comparison. Publicly available datasets with both single-cell and single-nucleus data that contain adipocytes provided an ideal test case. While mesothelial cells seems to be present in some single-cell sequencing datasets [26, 27], in these datasets we used both adipocytes and mesothelial cells are shown to be lost during single-cell sequencing but the nuclei are preserved and profiled in single-nucleus sequencing [28]. In our experiment, the cell-type proportion in pseudobulks was defined based on cell types observed in single-cell data, while the expression profiles themselves were derived from single-nucleus counterpart. Figure 4A shows the data origin for each component of the simulated data. This provided gold standard cell-type proportions with realistic missingness. The single-cell references are based on the single-cell profiles as well, which in this experiment has substantial depletion of two cell types, adipocytes, and mesothelial cells (see Methods for details; Supplemental Table 1 for cells and proportions present in each dataset). We then examined scenarios with and without added sources of noise. As in previous experiments, the residual is calculated with the deconvolution-calculated proportions and the single-cell reference. The residual is then factorized into 3 factors, from which one is expected to be correlated to the adipocyte’s and one to the mesothelial cell’s proportions. We observed variable results for deconvolution performance between scenarios where pseudobulks had noise added and those without as well as between experiments with random proportions and realistic proportions. Performance was overall lower in pseudobulks of realistic proportions when compared with those created using uniform random proportions for all deconvolution methods (Figure 4 B-D vs. Supplemental Figure 7). It is important to note that the two missing cell types encompassed over 50% of all cells in the mixtures (Supplemental Table 1). When comparing bulks with and without noise added, we also observed an expected decrease in deconvolution accuracy, as seen in Figure 4 B-D (left) vs. Supplemental Figure 7 (left panels of A-C).

**Figure 4.**
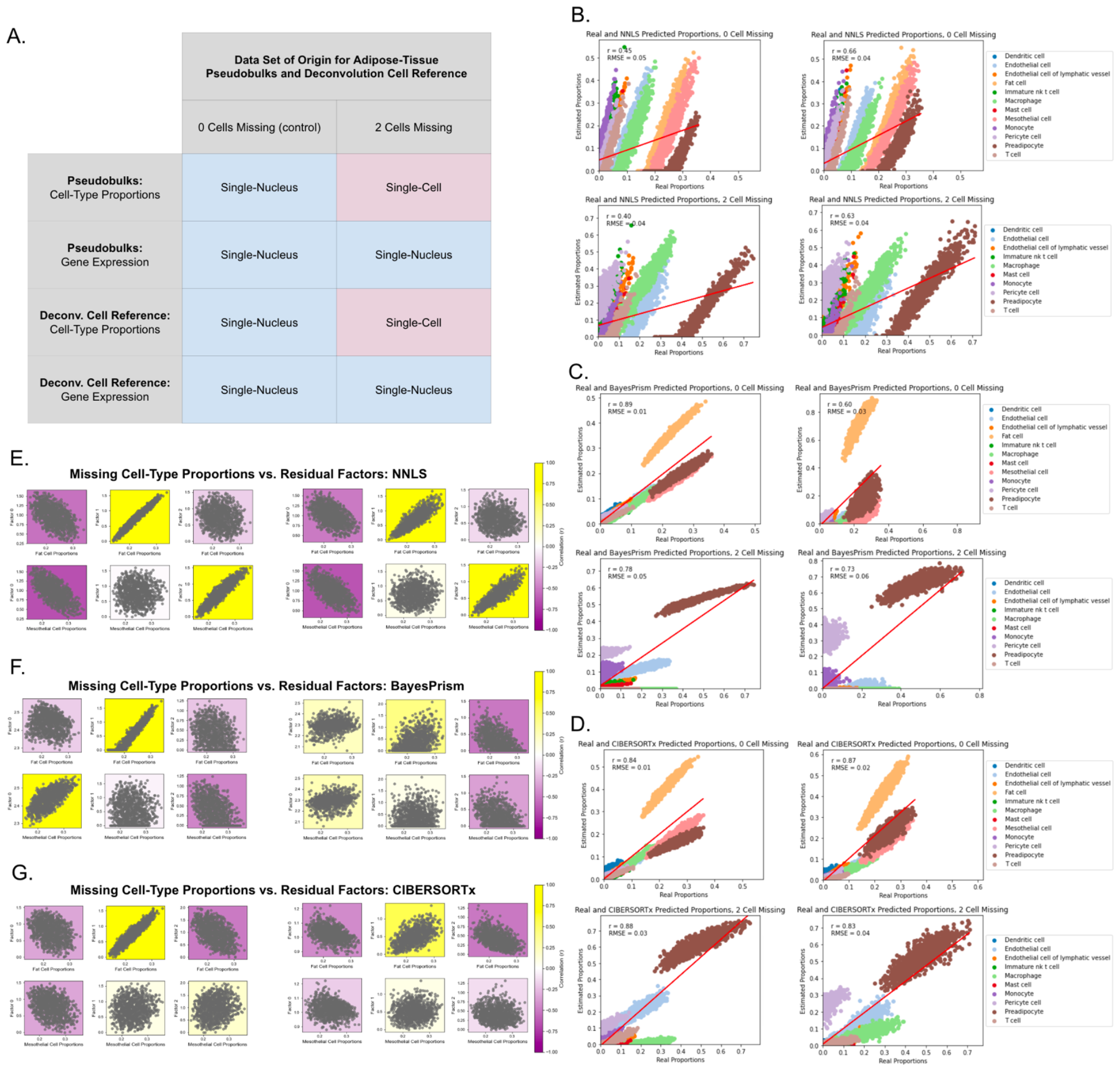
Single-nucleus and single-cell informed-pseudobulks comparing real and simulated (random) missing cell-type proportions. **A**. Pseudobulks created from single-nucleus RNA-seq adipose tissue cell expression, either with random proportions or with single-nucleus proportions (not missing any cell types). Single cell references for deconvolution are either with single-nucleus proportions (no cells missing, control), or using single-cell RNA-seq proportions (two cell types missing). Panels on the right (B-D) show real vs. calculated proportions for pseudobulks of realistic proportions in each of the deconvolution methods. The left column represents pseudobulks with no noise added, and the right column represents bulks with noise added. **B**. NNLS **C**. BayesPrism, and **D**. CIBERSORTx. The top plot of each panel represents the deconvolution with no cells missing (same cells as present in pseudobulks), and the bottom plot represents the proportions with two cells missing (no adipocytes or mesothelial cells). The red line in each plot represents the regression fit line. Each plot has RMSE and Pearson’s correlation noted. The bottom left panels show the correlation between the residual’s NMF factors and each of the missing cell-types’ proportions across pseudobulks for **E**. NNLS, **F**. BayesPrism and **G**. CIBERSORTx. Plots in the left column represent pseudobulks made with no noise added and those in the right column represent pseudobulks with noise added.

We then examined whether the residuals still contain some information on the missing cell types and found that pseudobulks with realistic proportions had lower correlation of missing cell-type proportions and residual’s factors when compared to random-proportioned pseudobulks (Figure 4 (E-G) vs. Supplemental Figure 7 (D-E)). However, we observed some correlation, especially in NNLS deconvolution (Figure 4E). When comparing pseudobulks with and without added noise, the noise lowered correlation further across deconvolution methods. Overall, this is not surprising given that implementing NMF in this manner assumes the residual contains no or minimal noise. In the presence of noise, NMF is expected to interpret the noise as part of the underlying factors, leading to inaccurate separation. When deconvolving noisy0pseudobulks, BayesPrism showed the lowest correlation (Figure 4 (E-G) left vs. right, and Supplemental Figure 7 (D-F) right vs. left) and highest RMSE values (Supplemental Figure 8) in the factor-proportion comparison. This is attributed to BayesPrism’s lower deconvolution accuracy with noise added.

### Recovery of missing cell-type’s expression in HGSOC bulk RNA-seq samples

We next sought to determine whether results consistent with this model would be observed in real-world data with bulk and matching single-cell data. We analyzed a dataset of 8 matched bulk and single-cell RNA-seq, for which we also have bulk RNA-seq counts of tissue that was dissociated before sequencing and thus lacking cells lost in dissociation (Dissociated bulks). For some samples, the location of the sample origin was available (Supplemental Figure 9A). We hypothesized that the Dissociated bulks were missing adipocytes when compared to typical non-dissociated bulks (Classic bulks). Our experimental design tested whether deconvolution of dissociated bulk (bulk and single-cell have same cell types) yielded a different residual than residuals of bulks with one extra cell type (adipocytes) when compared to their single-cell counterparts, and whether the missing cell’s expression was recoverable through the residual (Supplemental Figure 9B).

We first analyzed whether adipocyte-related genes were overall higher in residuals of classic bulks compared to dissociated bulks. Dissociated genes (characterized in the data’s original publication [24]) were much higher in dissociated bulks. As expected, CIBERSORTx barcode genes showed roughly even distribution, and adipocyte marker genes were higher in residuals of classic bulks (Figure 5 A-B). We also observed this in violin plots comparing the distributions of Classic and Dissociated residuals by gene group (Supplemental Figure 10).

**Figure 5.**
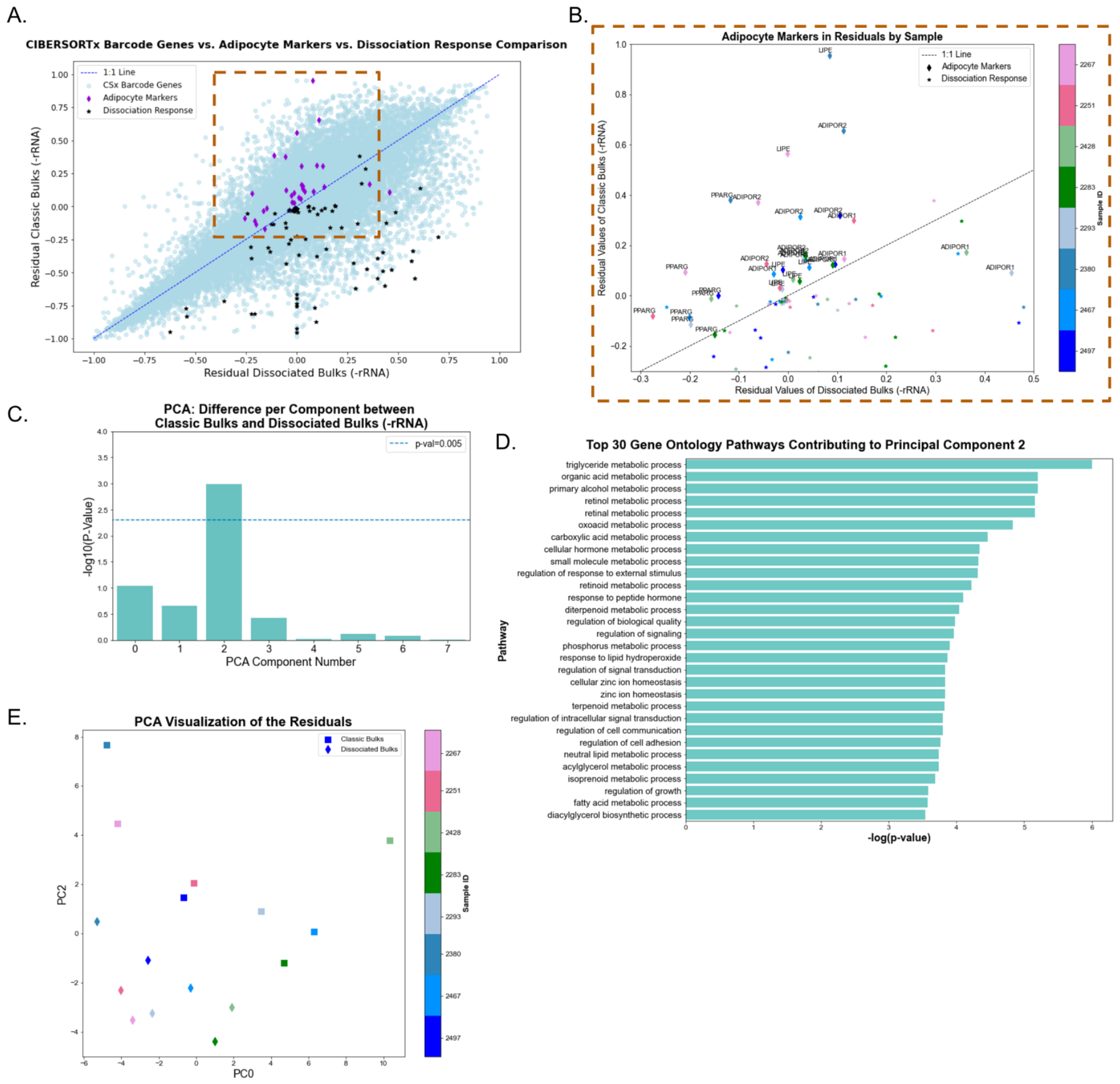
Residual analysis from HGSOC Dissociated and Classic bulk samples deconvolved with NNLS using matched single-cell RNA-seq data. **(A-B**) **Comparison of residual values of Dissociated Bulks (-rRNA) and Classic Bulks (-rRNA) at the gene and sample level**. Dissociated bulks residuals are on the x-axis, and Classic bulks are on the y-axis in both panels. CIBERSORTx barcode genes (calculated from all single-cell datasets), Adipose Markers, and Dissociation Response genes (from original paper) are compared. **A**. Scatter plot of gene groups of Dissociated bulks (x-axis) and Classic bulks (y-axis), with the 1 to 1 line in dashed blue. **B**. Zoomed-in version of panel A along the dotted line, coloring each gene by which sample it originates from. **(C-E) Factorization and Visualization of the Residuals of Classic Bulks (-rRNA) and Dissociated Bulks (-rRNA) combined. C**. Paired t-test results showing the negative log(p-values) for each of the means of the principal components. Components 0, 1, 3-7; all p-values > 0.05, and Component 2 showed a t-statistic of 4.41 and a p-value of 0.000595 **D**. Gene pathways from gene ontology analysis using GOrilla on the ordered genes contributing to Principal Component 2. **E**. PCA of each sample, classic and dissociated, within the PC0 and PC2 space.

We performed PCA on the residual from the analysis of both the Dissociated and Classic bulks (see Methods for details). Following the PCA, we conducted t-tests to assess whether the means of the principal components were statistically different between Classic and Dissociated bulks. We found that while most components showed no significant differences (PC 0, 1, 3-7; all p-values > 0.05), PC2 was significantly different with a t-statistic of 4.41 and a p-value of 0.000595, suggesting that this component captures a key variation attributable to the dissociation process (Figure 5C). We performed GOrilla [29] analysis of each PC to identify ontology terms associated with each PC (See Methods for details). This analysis revealed that for PC2, numerous pathways were associated with adipocyte function, including metabolism, lipid processing, and triglyceride biosynthesis (Figure 5D for PC 2, and Supplemental Figure 11 for the rest of the components). Few samples had available origin annotations, and we found the two omental samples were not the only ones with high PC2 levels. For example, sample 2428, derived from the ovary, was also high on the PC2 axis (Figure 5E). While the omentum is known to be a tissue rich in adipocytes, other tissues can also have substantial adipocyte content, making a 1:1 correspondence between high-adipose and originating site for the sample difficult to draw.

To evaluate a method other than PCA, we also factorized the Dissociated and Classic bulks using NMF (Supplemental Figure 12). We again used GOrilla [29] on the ordered list of genes contributing to each factor. Although there were many statistically significant pathways, we found no component related to adipocytes through this factorization method (Sup. Figure 12 B-E). This may be because, unlike PCA, NMF does not enforce orthogonality between factor matrices, making highly correlated factors difficult to disentangle, which might be the case for the adipocyte signal in the residuals. Taken together, while our results demonstrate information present in the residuals can reveal missing cell types, the ability to extract and identify those depends on the dataset, deconvolution method, and analytical approach applied to the residuals.

## Discussion

The goal of this paper is not to present a novel method but to raise awareness about the intricacies of RNA-seq technologies. Our study aimed to explore the deconvolution performance of NNLS, BayesPrism, and CIBERSORTx under varying conditions of missing cell types and realistic proportions using both simulated pseudobulks and experimental bulk RNA-seq datasets. We describe the impact of missing cell types, their similarity, and having bulks with realistic proportions on the deconvolution accuracy.

In our experiments with distinct immune cell types, we observed a decrease in deconvolution performance as the number of missing cell types increased, similarly to previous work done with one cell type missing [11, 20]. We also observed the missing cell-types’ proportions have very high correlation to one of the residual’s factors.

When applying the deconvolution methods to the PBMC3k dataset, we observed that the cell-type’s similarity plays a crucial role in whether we can recover the missing cell-types’ proportions through NMF. The more similar the cell type is to others, the more that cell type’s corresponding NMF factor is convolved with others in all three deconvolution methods (Figure 2). This is in agreement to previous work [22].

Furthermore, we found that deconvolution performance was consistent regardless of the expression similarity among missing cell types. We also found comparable performance across NNLS, BayesPrism, and CIBERSORTx in recovering missing cell-types’ proportions. However, the NNLS residual factor had higher correlation overall, which we attribute to NNLS being a regression model, and our methodology of calculating the residual is based on regression’s residuals.

The impact of realistic proportions and simulated noise on deconvolution performance was explored in the context of adipose tissue datasets. Realistic proportions of adipocytes and mesothelial cells, which collectively constituted a significant portion of the mixtures, led to a decrease in deconvolution accuracy across all methods. The addition of noise further diminished the correlation between the missing cell types’ proportions and residual factors. Despite this, the residuals still contained information about the missing cell types, emphasizing the potential utility of our approach even under challenging conditions.

Applying our approach to data generated to understand HGSOC cell-type proportions suggests that such datasets exhibit signatures of these missing cell types. While no gold standard is available in this context, gene ontology analysis of the residuals of Dissociated bulks and Classic bulks shows that adipocyte-related gene sets are observable through PCA. The results also enable hypothesis generation about samples with incomplete metadata. For example, of the 4 samples of unknown origin, we hypothesize sample 2380 is also of omental origin or otherwise contains many adipose cells. This analysis could be a starting point for future methods seeking to uncover missing cell-type identities within bulk RNA-seq data.

In conclusion, we examined the impact of missing or lost cell types on deconvolution methods, emphasizing the importance of understanding these constraints for meaningful biological insights. As researchers delve into increasingly complex tissues, these types of factors can influence results and estimates of tissue composition. Recognizing these constraints not only refines our interpretation of results but also paves the way for continued advancements in technology and methodologies. The presence of missing cell-type’s signal in the residual suggests multiple paths for future methods; while the ideal reference is likely matched from the same participants, perhaps residuals could enable searching of cell-type reference libraries for profiles to augment deconvolution. Alternatively, iterative procedures could be used to estimate missing cell types, refine deconvolution, and potentially repeat the process. In either case, acknowledging and addressing these challenges will undoubtedly enhance the robustness and reliability of our findings, fostering a more profound and accurate understanding of the biological intricacies encoded in RNA-seq data.

## Methods

### Data

We used three datasets for this study, all of which are available online as count matrices through NCBI GEO. Each was processed with adjustments for each data type, outlined in Supplemental Table 2, along with cell-type assignments, data links, and related figures. The cell types were assigned according to each data set’s original study, apart from the 10x Genomics PBMC3k dataset, for which we used Scanpy’s [30] tutorial (https://scanpy-tutorials.readthedocs.io/en/latest/pbmc3k.html) cell assignments.

We first used single-nucleus white adipose tissue RNA-seq data and extracted the expression of 5 cell types (macrophage, T cell, B cell, monocyte, dendritic cell). We then expanded in complexity and used a PBMC3k single-cell RNA-seq dataset from 10x Genomics. We also used the single-cell and single-nucleus dataset of adipose tissue with real missing cell types in the single-cell counterparts compared to the single-cell. Cell-type similarities vary between each dataset, and these are shown in a correlation distance dendrogram in Supplemental Figure 5. We created simulated bulks with each of these datasets.

We also leveraged HGSOC datasets from a previous study [17] from our group that contains data on 8 tumors, with matched single-cell RNA-seq, classic bulk RNA-seq with ribosomal RNA depletion (Classic bulk - rRNA), dissociated cells that were bulk RNA-sequenced with ribosomal RNA depletion (Dissociated bulk -rRNA) and dissociated cells that were also bulk RNA-sequenced with poly-A tail capture (Dissociated-bulk polyA). Our dataset consists of 2 known omental samples and 2 ovarian samples, and 4 samples with unknown origin.

### Pseudobulks for deconvolution of known cell-type proportions

Pseudobulks were generated using single-cell or single-nucleus data from the sources detailed in Data section, apart from the HGSOC data, for which we have matched real bulks. We created pseudobulks with both random proportions, and realistic proportions. The proportions for each pseudobulk with random proportions are defined as a vector of random numbers for random proportions, enforcing a sum-to-one constraint. For the realistic proportions, we used a vector of the real cell proportions in the data, with added Gaussian noise for variability in each cell-type’s proportions for all bulks.

For each pseudobulk, we defined the total number of cells to be 5,000. We multiplied the proportion vector (described above), defined as either random or realistic, by the number of cells per bulk (5000) to get the number of cells, per cell type, in each bulk. Cells were then sampled from the original dataset to match this number, representing the number of cells per bulk. We summed the expression of the sampled cells for each gene, giving the final pseudobulk count.

In the experiments using real missing cell types in adipose tissue, we also have single-nucleus and single-cell data. The pseudobulks were created by using the expression of single-nucleus cells (no cells missing). For the pseudobulks with noise added used in this experiment, we added Gaussian noise to simulate library size and general variability noise seen in real bulk RNA-seq. We end up with four types of pseudobulks (random with no noise, random with noise, realistic with no noise, realistic with noise), all based on single-nucleus cells’ expression.

### Cell reference for deconvolution

The single-cell references were created using the same single-cell dataset as the pseudobulks, containing either the same cell types as the pseudobulks, or with missing cell types. The single-cell reference for deconvolution was constructed by sampling with replacement from each cell type to reach a uniform count of 10,000 cells per cell type to standardize the data and reduce potential bias caused by varying cell counts across types. We then added these counts per cell type, which gave us a signature expression per cell type, per gene. We used this reference as having 0 missing cell types, and subsequently iteratively depleted the reference of cumulative randomly chosen (or specified) cell types. We removed cells until the reference contained a minimum of 2 present cell types. The pseudobulks of five distinct cell types from single-nucleus adipose tissue were deconvolved with 0, 1, 2, and 3 missing cell types. The PBMC3k pseudobulks were deconvolved with 0,1, 2, 3 and 4 missing cell types, both with and without similar expression. For the single-cell/single-nucleus adipose tissue, we only considered 0 missing cell types (real single-nucleus proportions) or 2 missing cell types (real single-cell proportions). This is shown in Figure 5. For each experiment, we end with a distinct single-cell reference matrix with a specific number of known missing cell types compared to the pseudobulks.

For NNLS, the single-cell reference and each pseudobulk are kept in the linear scale to improve deconvolution performance, as observed by a benchmarking study [20]. We MinMax scaled the pseudobulks and the reference prior to deconvolution. Prior to MinMax scaling, we clipped the top 95% highest values from the matrix, so that gene expression from high-expressing genes would not skew our scaled values. The pseudobulks were not scaled for BayesPrism and CIBERSORTx, in accordance with the metholodgy’s specifications. The NNLS reference was filtered to only contain the barcode genes computed through CIBERSORTx reference matrix with 0 cells missing (containing all cells’ barcode genes) to maximize each cell’s signal. BayesPrism and CIBERSORTx references contained all genes of their respective references as computed by the deconvolution method.

For the adipose single-cell/single-nucleus experiment (Figure 4 and Supplemental Fig. 7), single-nucleus data was used as the reference (with no missing cell types) since it contains all cells and matched the expression of the pseudobulks. For the reference with missing cell types (2 in this real case), we created the reference with the single-cell proportions (missing adipocytes and mesothelial cells) but sampling the expression of each cell from the single-nucleus data. In the case of real HGSOC bulk deconvolution, we combined the single-cell matched datasets from the bulks into one reference used to deconvolve all the bulks to maximize the number of cells present.

The adipose single-nucleus/single-cell dataset contains cells from multiple patients, making it too large for all cells to be reasonably used in deconvolution for CIBERSORTx and BayesPrism which create a single-cell reference based on each cell. BayesPrism literature shows robust performance with references containing between 100 to 500 cells [11], and CIBERSORTx [12] only tested performance up to 500 cells per cell type.

Therefore, we limited the number of cells to 2000 for these deconvolution methods when using these large data without compromising performance.

### Deconvolution (NNLS, CIBERSORTx, BayesPrism)

We used NNLS (regression-based), CIBERSORTx (marker-gene based) and BayesPrism (Bayesian-based) deconvolution in both pseudobulks with ground truth and real bulks. Software versions for each deconvolution method used are NNLS v.1.13.0, BayesPrism v2.2, CIBERSORTx v1.05.

For CIBERSORTx and BayesPrism, we followed the specifications of the methodology for deconvolution. All methods were given a reference with no cell types missing and with cell types missing as outlined in Cell reference for deconvolution section, and our pseudobulk with all cell types to be deconvolved. This process was repeated for 1000 pseudobulks, with references with no missing cell types and with missing cell types. The NNLS deconvolution was run with SciPy’s NNLS’s algorithm [25] with default settings.

BayesPrism was run through InstaPrism [31], and CIBERSORTx was run through the Docker container with a token from the CIBERSORTx team.

### Residual calculation

Residuals were calculated by subtracting the recreated bulks from our original input pseudobulks matrix, as shown in Figure 1B. The recreated bulks matrix is computed by multiplying our deconvolution-calculated proportions by the single-cell reference (MinMax scaled) used in the deconvolution method. This residual matrix (samples by genes) is then calculated by subtracting the recreated bulks from our original pseudobulks. This residual represents the deviation from the expected proportions based on the deconvolution results.

CIBERSORTx uses a small number of marker genes for each cell type, called “barcode genes”, so the single-cell reference matrix created contains only a sample of the genes used. To be able to calculate a comparable residual, we used the reference as computed for NNLS for the recreated matrix instead, but we multiplied with estimated proportions from CIBERSORTx.

### Residual factorization and evaluation

The residual matrices contain pseudobulks (rows) by genes (columns) for each reference used in deconvolution (with all cell types and with missing cell types). Because the residuals have negative values, we shift the distribution of values by the minimum value needed for all values to be greater than 0. We factorize each residual using non-negative matrix factorization (NMF) [25] with Nonnegative Double Singular Value Decomposition (nnsvd) initiation, with the number of components set to be equal to the number of missing cell types plus one, and with maximum iterations of 10000 for each. Each NMF components’ values are scaled to be between 0 and 1 (the scale of proportions), and further compared to each of the missing cell type’s proportions for only pseudobulks in which we expect the distribution of proportions to be distributed between nearly 0 and 1 (random proportions). For realistic proportions, we do not expect the proportions to have that distribution, and therefore the RMSE is not meaningful and not calculated in those cases. For random proportions, the RMSE and Pearson’s correlation for each combination of component missing cell type is computed and recorded. For realistic proportions, only Pearson’s correlation is recorded.

### Evaluation of residuals in high-grade serous ovarian cancer data

We used the real Classic (-rRNA) or Dissociated (-rRNA and polyA) bulks for deconvolution using NNLS, (experimental design, Supplemental Figure 1). The Dissociated polyA bulks were used as a control, since the polyA tail capture protocol and the ribosomal depletion protocol might cause specific changes in the bulk data that we have not characterized in this work. Therefore, Classic bulks and Dissociated bulks, both with -rRNA protocols, were compared. The residuals were calculated as in previous sections of this paper for Dissociated and Classic (-rRNA) and Dissociated (polyA).

In contrast to our previous experiments, no ground truth was available for the dataset of real bulks, so we could not compare proportions to the residual matrix’s factors using the previous strategy. Instead, we compared the residual values for each bulk type and used a non-parametric Wilcoxon t-test to compare the distributions.

We also performed factorization of the combined Classic and Dissociated (both -rRNA protocols) residuals with PCA and NMF to evaluate whether the residual would show information of the hypothesized missing adipocytes in a PCA or NMF component. The PCA was implemented using the PCA function from the Scikit-learn [32] library (v.1.4.2). Scatter plots were generated to visualize the distribution of the two sample types across the principal components. The plots were colored by sample type to facilitate visual differentiation. A two-sample t-test was conducted to identify significant differences between the Classic and Dissociated bulks across the principal components. The analysis was performed for each principal component separately, and p-values were adjusted using the Bonferroni correction to control for multiple comparisons.

For each of the eight principal components, genes were ranked based on their loadings, calculated as the product of the PCA components and the square root of the explained variance. The genes contributing the most to each component were identified and listed in order. This gene list was later used for further biological interpretation by conducting gene ontology analysis using GOrilla [29], a tool for discovering enriched GO terms in ranked lists of genes. The genes from each principal component were input as a single ranked list, focusing on biological processes. The top 30 significant pathways, based on the lowest p-values, were extracted and plotted to illustrate the variation most associated with the genes contributing to the principal components. The significance of the pathways was visualized through bar plots, displaying the negative logarithm of the p-value for each pathway associated with the principal components.

Similarly, NMF was applied to the same residual data with the same software and parameters outlined in the section on residual factorization and evaluation above. A two-sample t-test was conducted for each principal and NMF component to identify significant differences between the classic and dissociated bulks. The analysis was performed for each component separately, and p-values were adjusted using the Bonferroni correction to control for multiple comparisons. Gene ontology analysis was also performed using GOrilla [29], and the top 30 significant pathways, based on the lowest p-values, were extracted, and plotted to illustrate the processes most associated with the genes contributing to the NMF factors.

## Code availability

The code developed for this study is available at our GitHub repository: https://github.com/greenelab/pred_missing_celltypes under BSD 3-Clause License. The repository includes all necessary scripts and a README file with setup instructions and usage guidelines. For updates or assistance, users can refer to the repository or open an issue for queries. Our aim is to support transparency and reproducibility in computational research through this open-access resource. We’ll create an archival snapshot of the repository on Zenodo or Figshare once we have a version of code associated with an accepted paper.

