## Supplemental Figures for "Missing cell types in single-cell references impact deconvolution of bulk data but are detectable"

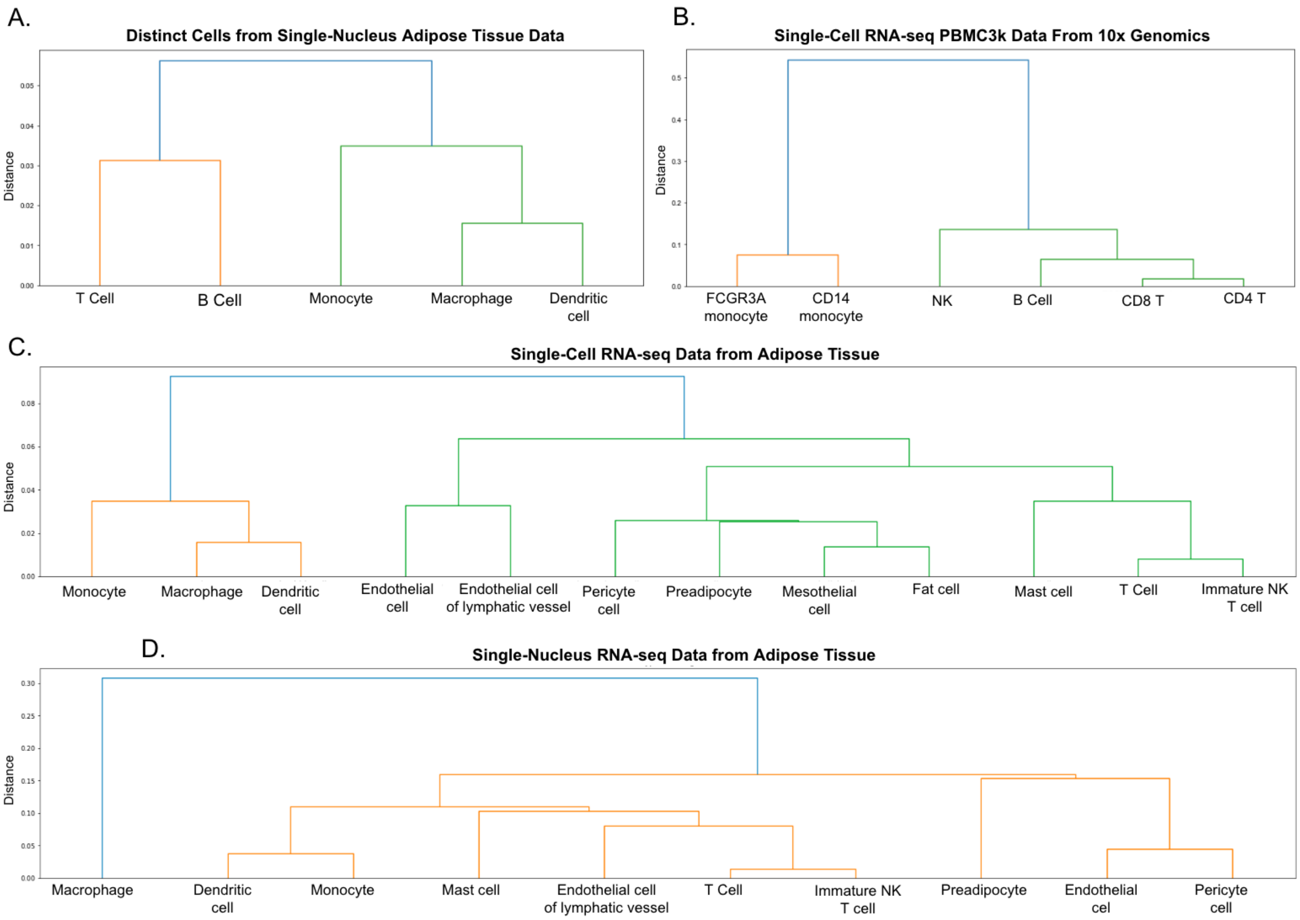


***Supplemental Figure 1. Correlation distance dendrogram between all cell types in each of the datasets used, representing differences in gene expression by cell type****. Distance is calculated as the inverse of Pearson's correlation with each cell-type's expression. The greater the distance, the more different two cell types are.* ***A.*** *Dendrogram of 5 curated cell types extracted from single-nucleus adipose tissue dataset (shown in C).* ***B.*** *Dendrogram of all cell types extracted from PBMC3k 10x Genomics dataset.*

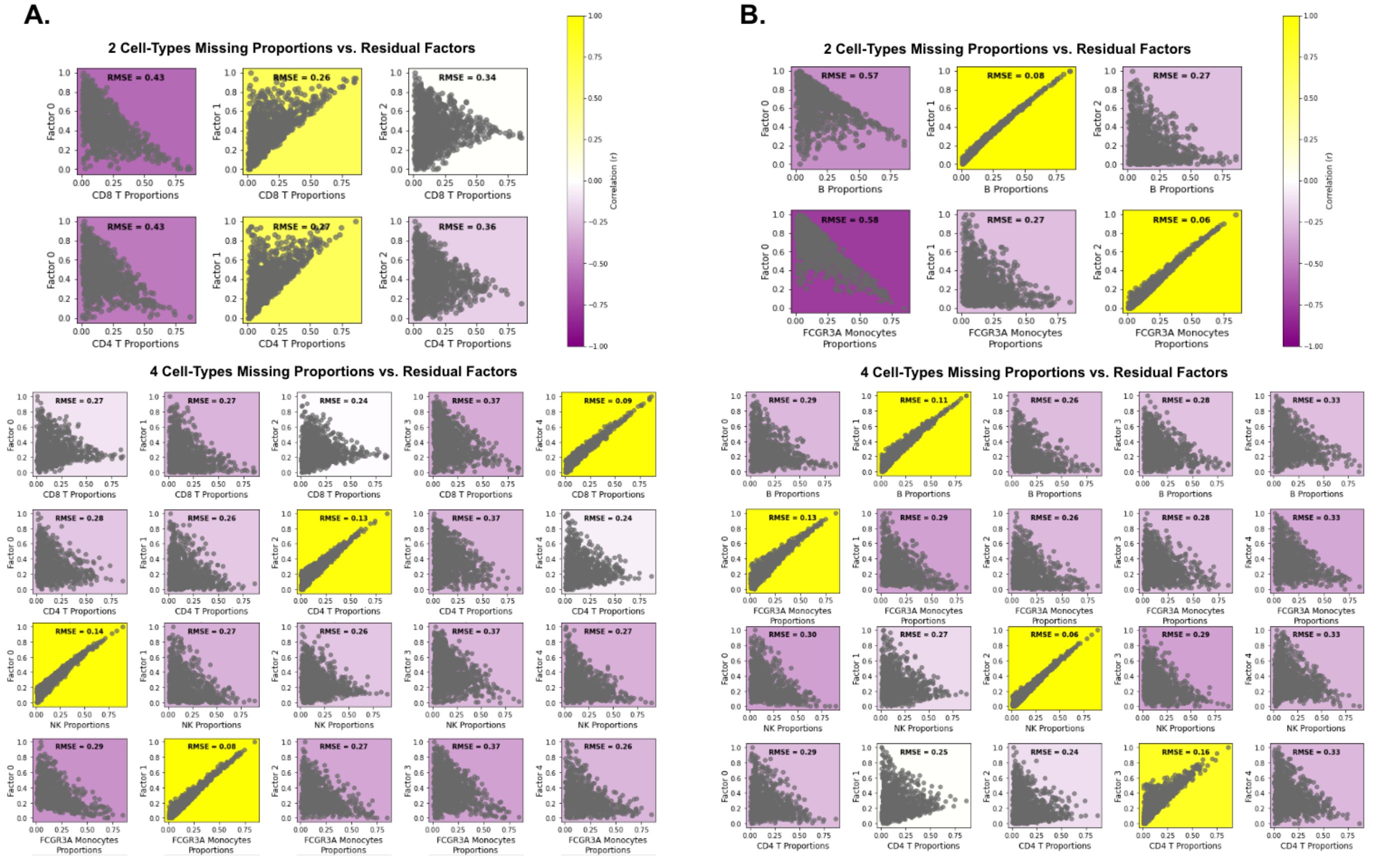


**Supplemental Figure 2. *NNLS Deconvolution of PBMC3k Pseudobulks with Random Proportions*:** *A portion of these data is shown in Figure 3 on the main paper. We remove one, two, three and four cell types from the deconvolution reference. These cell types are selected to have low correlation in gene expression* ***A.*** *or selected randomly from cells with similar expression.* ***B.*** *The residual matrix is calculated and factorized with NMF. Each factor is then correlated to each of the missing cell-type's proportions. Pearson's correlation (color bar) is shown in the coloring of each plot, and the RMSE value between the residual factor and the cell-type proportions are noted. Note: The rest of the data (for one and 3 missing cell types) is in Figure 2 of main text.*

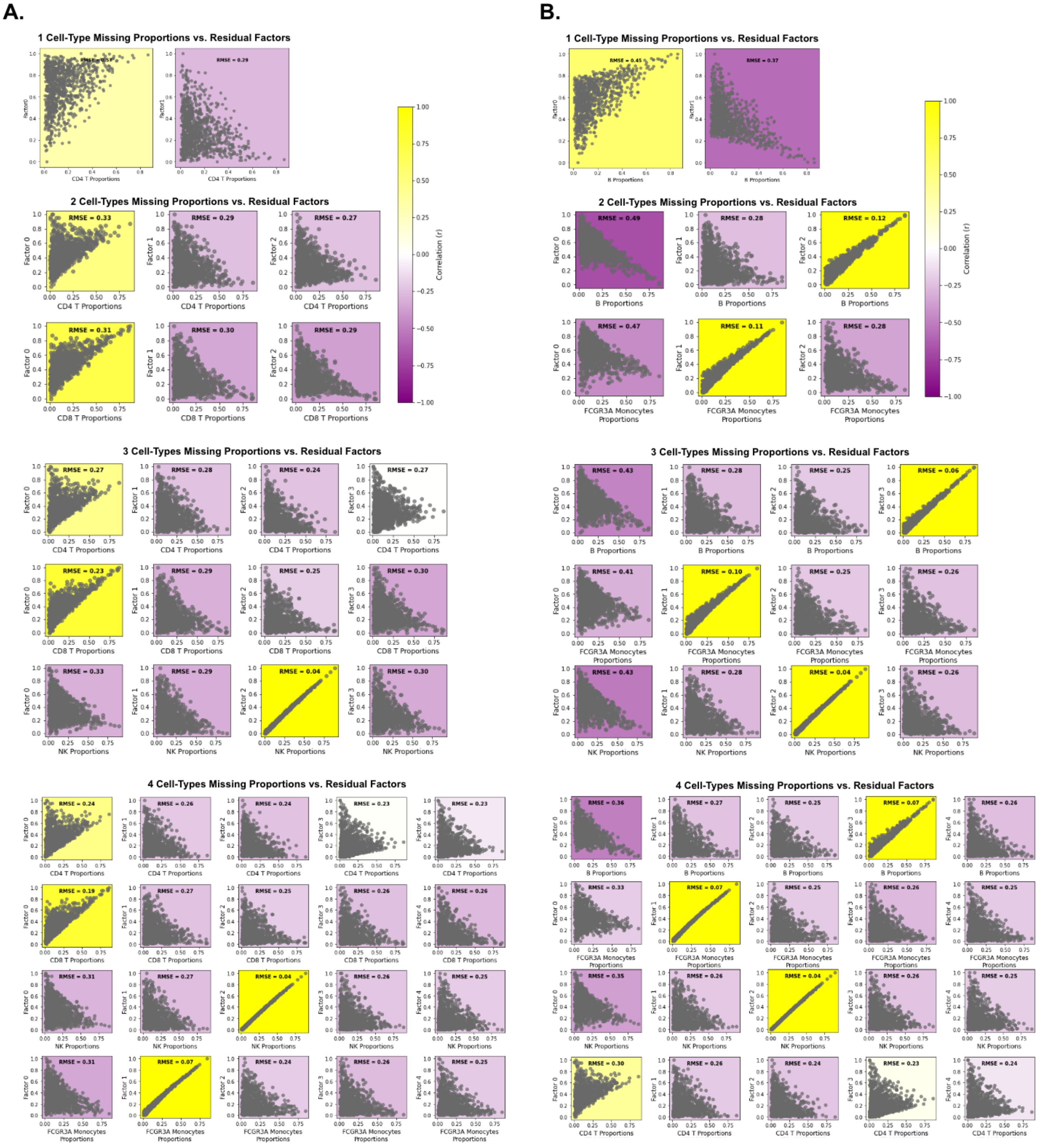


***Supplemental Figure 3. BayesPrism Deconvolution of PBMC3k Pseudobulks with Random Proportions.*** *We remove 1, 2, 3 and 4 cell types from the deconvolution reference. These cell types are selected to have low correlation in gene expression.* ***A.*** *or randomly selected cells having similar expression.* ***B.*** *The residual matrix is calculated and factorized with NMF. Each factor is then correlated to each of the missing cell-type's proportions. Pearson's correlation (color bar) is shown in the coloring of each plot, and the RMSE value between the residual factor and the cell-type proportions are noted.*

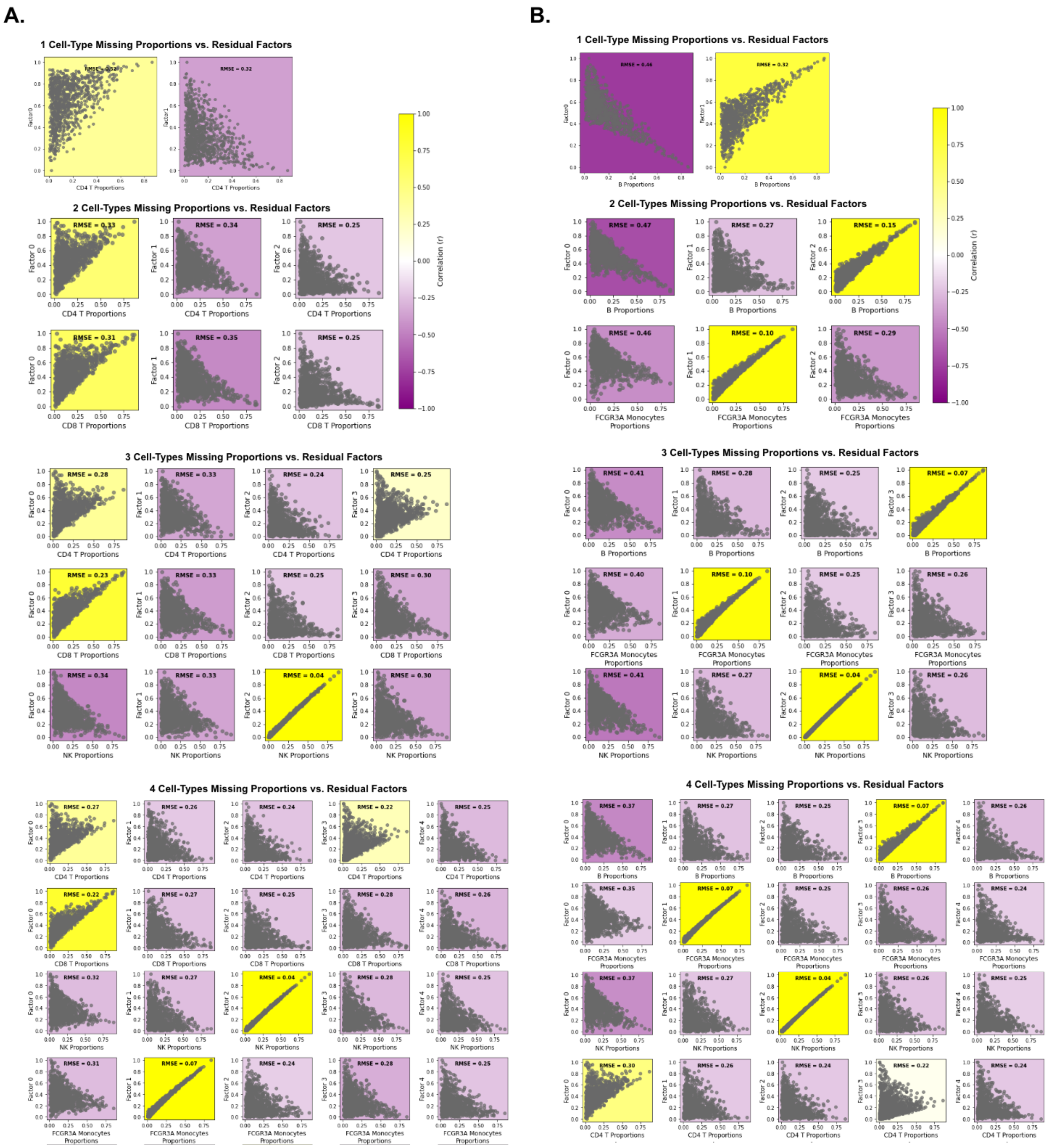


***Supplemental Figure 4. CIBERSORTx Deconvolution of PBMC3k Pseudobulks with Random Proportions:*** *We remove 1, 2, 3 and 4 cell types from the deconvolution reference. These cell types are selected to have low correlation in gene expression.* ***A.*** *or randomly selected cells having similar expression.* ***B.*** *The residual matrix is calculated and factorized with NMF. Each factor is then correlated to each of the missing cell-type's proportions. Pearson's correlation (color bar) is shown in the coloring of each plot, and the RMSE value between the residual factor and the cell-type proportions are noted.*

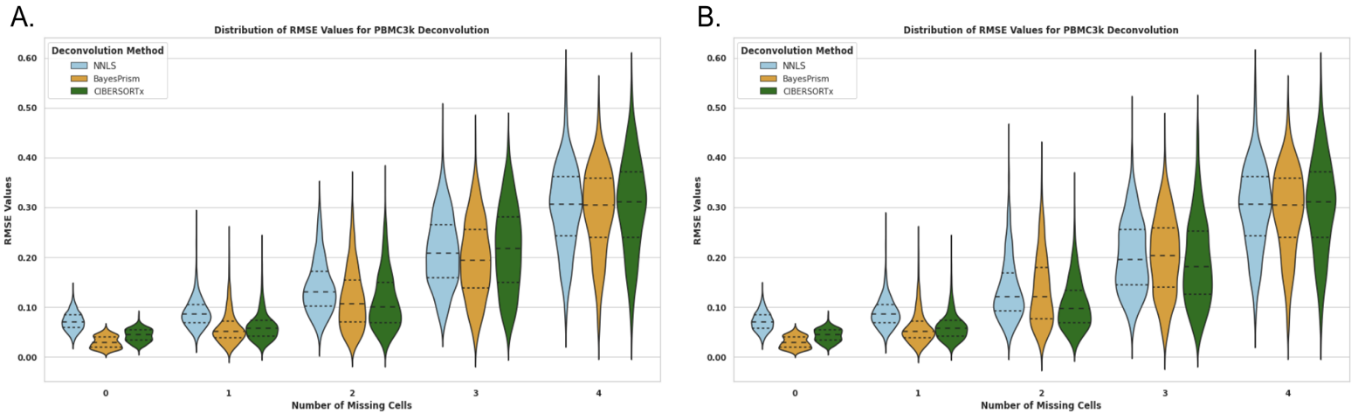


***Supplemental Figure 5. Comparison of deconvolution performance in NNLS, BayesPrism and CIBERSORTx for PBMC3k pseudobulks across number of missing cell types. A.*** *Violin plots showing deconvolution performance when deleting cell types that are not highly correlated.* ***B.*** *Violin plots showing deconvolution performance when deleting cell types that are highly correlated. Y axis represents RMSE value of calculated proportions vs. real pseudobulk proportions. Non-correlated cell types have only slightly better performance than when correlated cell types are removed.*

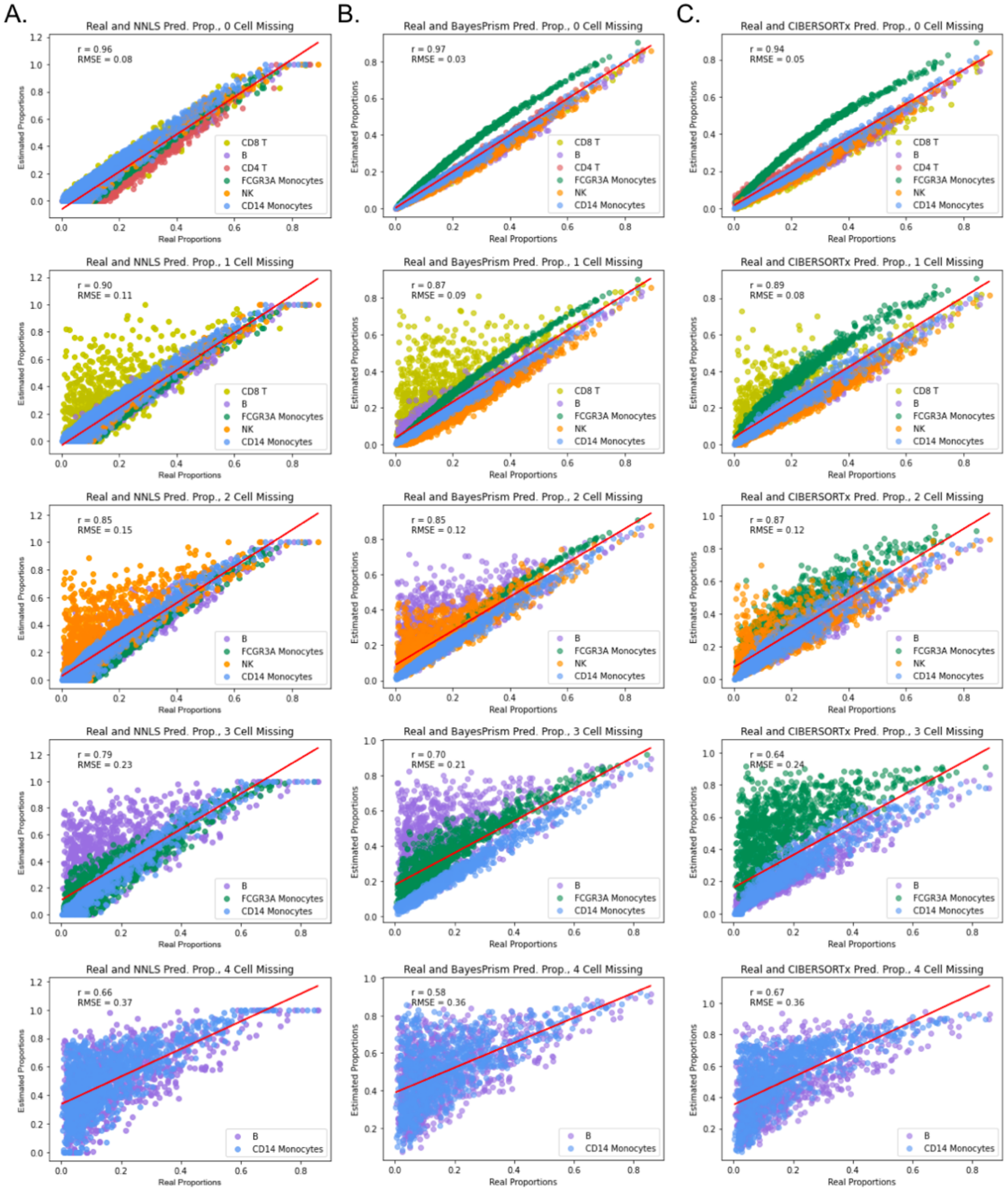


***Supplemental Figure 6. Correlated cell types being removed does not impact the performance of the deconvolution methods substantially.*** *From left to right,* ***A.*** *NNLS,* ***B.*** *BayesPrism and* ***C.*** *CIBERSORTx performance (real vs. estimated) proportions. First row corresponds to the control (0 missing cell types), second row corresponds to 1 missing cell type, then 2 missing cell types, up to 4 missing cell types in the last row. Pearson's correlation and RMSE values are noted in each plot.*

**A.** **Single-Nucleus Cell Content (0 missing)**

| Cell Type | No. Cells | Proportion |
|---|---|---|
| **Fat cell** | **24825** | **0.2197** |
| **Mesothelial cell** | **26276** | **0.2324** |
| Dendritic cell | 679 | 0.0060 |
| Endothelial cell | 11480 | 0.101578 |
| Endothelial cell of lymphatic vessel | 2339 | 0.020696 |
| Immature NK T cell | 1073 | 0.0095 |
| Macrophage | 13625 | 0.1206 |
| Mast cell | 883 | 0.0078 |
| Monocyte | 709 | 0.0063 |
| Pericyte cell | 1165 | 0.010308 |
| Preadipocyte | 26941 | 0.2384 |
| T cell | 3022 | 0.026739 |
| **Total** | **113017** | **1.00000** |

**B.** **Single-Cell Cell Content  (1 missing)**

| Cell Type | No. Cells | Proportion |
|---|---|---|
| Dendritic cell | 983 | 0.053164 |
| Endothelial cell | 542 | 0.029313 |
| Endothelial cell of lymphatic vessel | 157 | 0.008491 |
| Immature NK T cell | 389 | 0.021038 |
| Macrophage | 1410 | 0.076257 |
| Mast cell | 53 | 0.002866 |
| Monocyte | 636 | 0.034397 |
| Pericyte cell | 52 | 0.002812 |
| Preadipocyte | 13461 | 0.728015 |
| T cell | 807 | 0.043645 |
| **Total** | **18490** | **1.000000** |

***Supplemental Table 1**. **Cell content of both adipose tissue datasets.*** *Single-Nucleus* ***A.*** *and Single-Cell* ***B.*** *RNA-seq cell types, number of cells, and proportions. The single-nucleus proportions are used to create the realistic-proportioned pseudobulks, and the single-cell proportions are used to create the cell reference for deconvolution. Cell types that are missing in single-cell are colored in blue.*

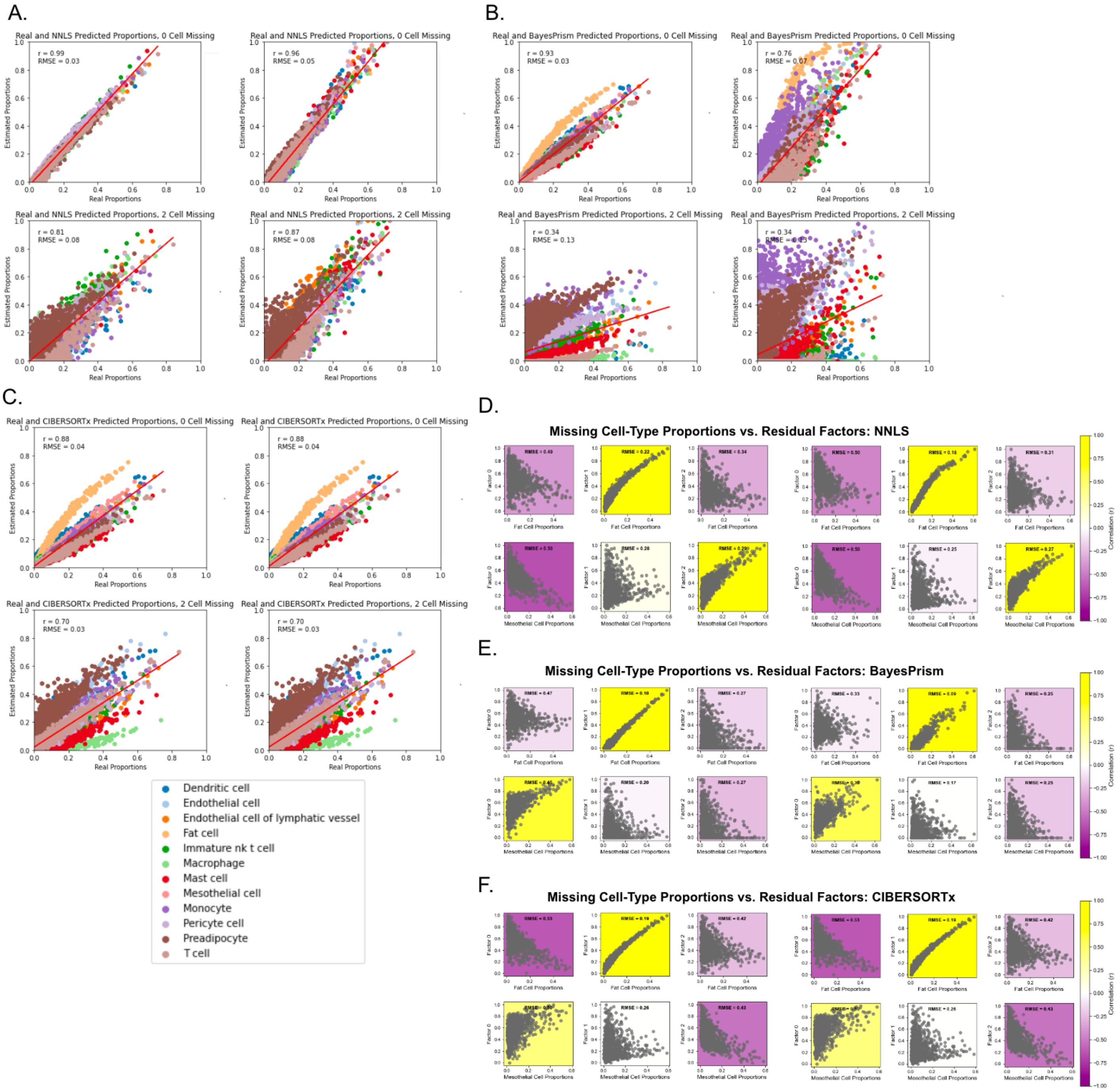


***Supplemental Figure 7. The left panel (A-C) shows the real vs. calculated proportions for pseudobulks of realistic proportions in each of the deconvolution methods.*** *The left column of these panels represents pseudobulks with no noise added, and the right column represents bulks with noise added.* ***A.*** *NNLS with no noise and NNLS with noise,* ***B.*** *BayesPrism with no noise and BayesPrism with noise,* ***C.*** *CIBERSORTx with no noise and CIBERSORTx with noise. The top panel of each represents the deconvolution with no cells missing (same cells as present in pseudobulks), and the bottom plot represents the proportions with 2 cells missing (no adipocytes or mesothelial cells), as seen in single-cell RNA-seq. The red line in each plot represents the regression fit line. Each plot has RMSE and Pearson's correlation noted.* ***The panels on the left (D-F) show the residual's factors of pseudobulks with random proportions compared to missing cell-type's proportions.*** *Plots on the left represent pseudobulks with no noise, and panels on the right represent pseudobulks with noise, each deconvolved with:* ***D.*** *NNLS,* ***E.*** *BayesPrism, and* ***F.*** *CIBERSORTx.*

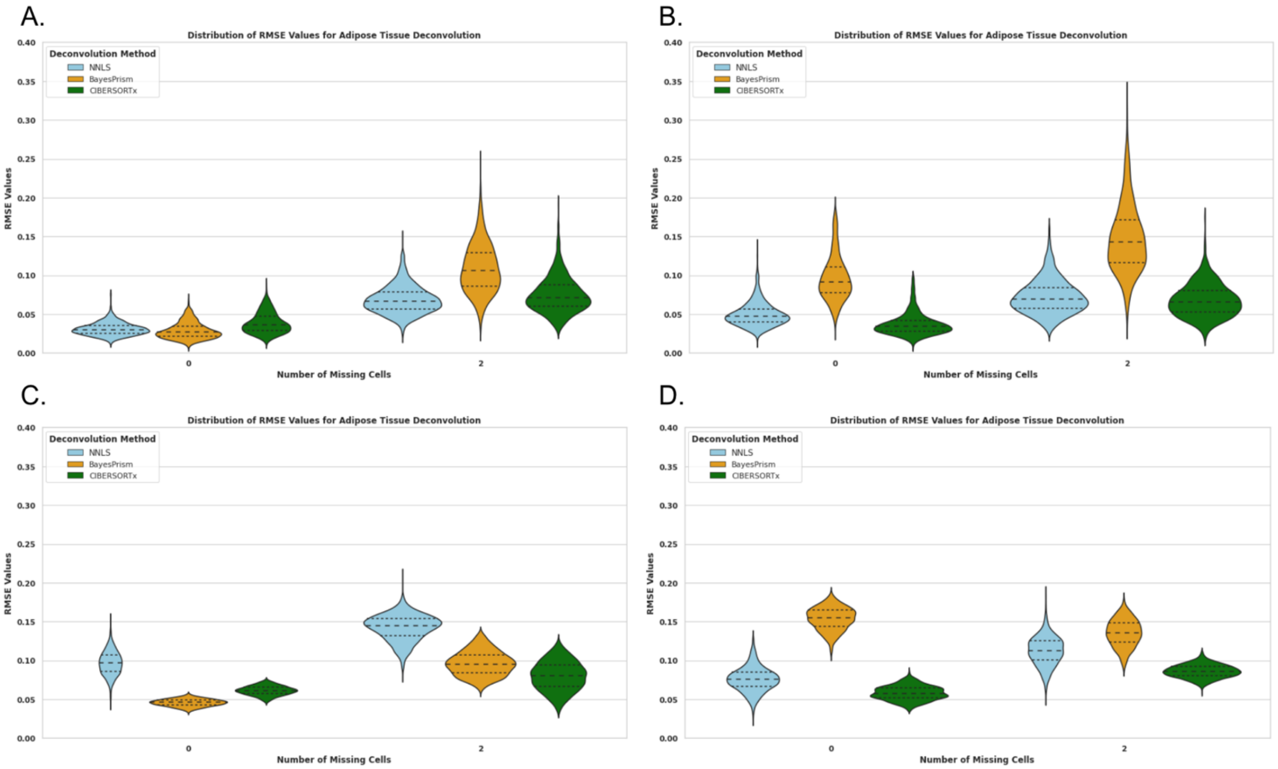


***Supplementary Figure 8. Calculated RMSE for each deconvolution method**. Panels on the right represent pseudobulks with noise, and panels on the left represent pseudobulks without noise. The top panels represent pseudobulks with random proportions, and the bottom panels represent pseudobulks with realistic proportions. **A. No noise, random, B. Noise, random, C. No noise, realistic, D. Noise, realistic.***

A.

| Sample Number | Tissue of Origin |
|---|---|
| 2267 | Omentum |
| 2251 | Omentum |
| 2428 | Ovary |
| 2283 | Right Ovary |
| 2293 | Unknown |
| 2380 | Unknown |
| 2467 | Unknown |
| 2497 | Unknown |

B.

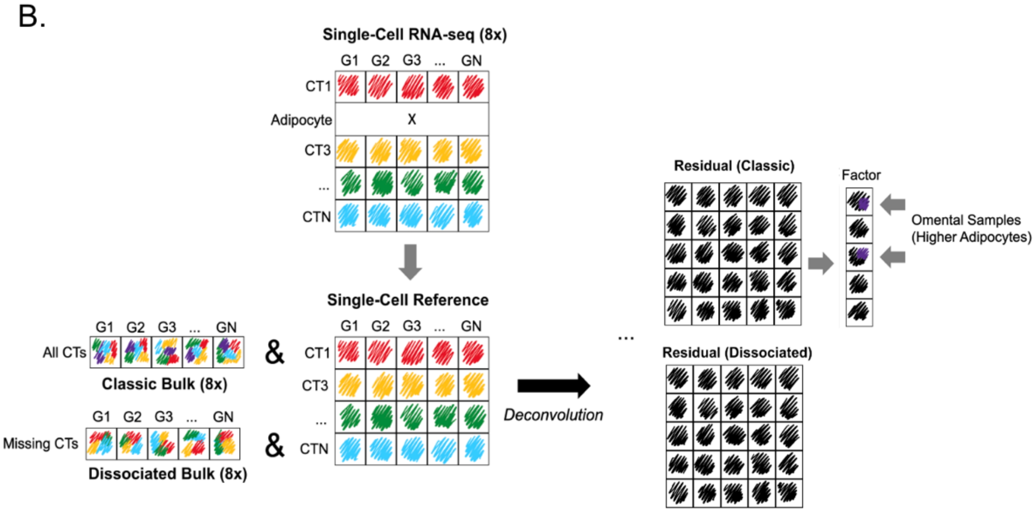


***Supplemental Figure 9. Analysis of experimental (not simulated) classic and dissociated bulks deconvolved with matched single-cell RNA-seq data. A.*** *Table showing the tissue of origin for each sample.* ***B.*** *Schematic illustration of experimental design. The dissociated bulks are hypothesized to match the cell types in the single-cell data, therefore are considered to have no cell types missing. The classic bulks are hypothesized to have at least one cell type missing (adipocytes). The residual is calculated as previously, and one of the residual's factors is expected to match adipocyte proportions.*

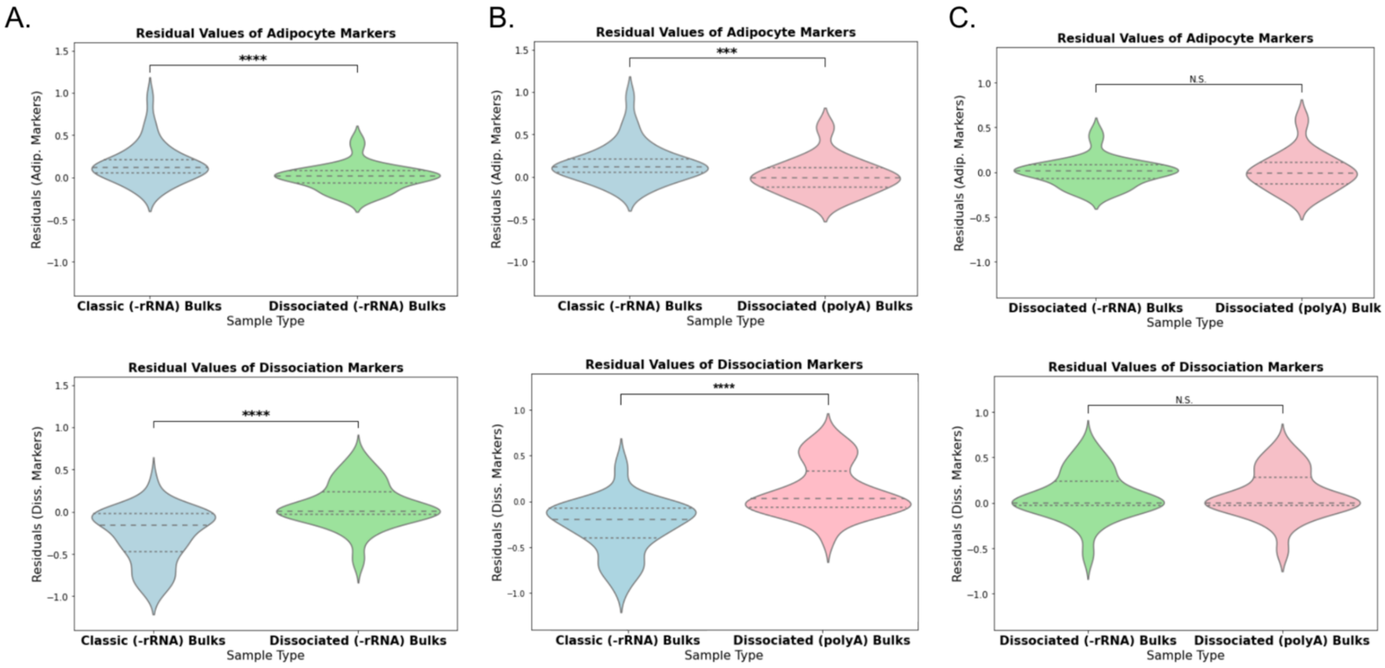


***Supplemental Figure 10. Violin plots of the Residual values in Adipocyte Markers (top), and Dissociation Response Markers (bottom). A.*** *Compares the adipocyte and dissociation values in the residuals of Classic and Dissociated Bulks (both -rRNA).* ***B.*** *Compares the adipocyte and dissociation values in the residuals of both Dissociated Bulks (polyA and -rRNA).* ***C.*** *Compares the adipocyte and dissociation values in the residuals of Classic Bulks (-rRNA) and Dissociated Bulks (polyA). Asterisks in all plots mark the statistical significance of p-values computed with Wilcoxon T-Test.*

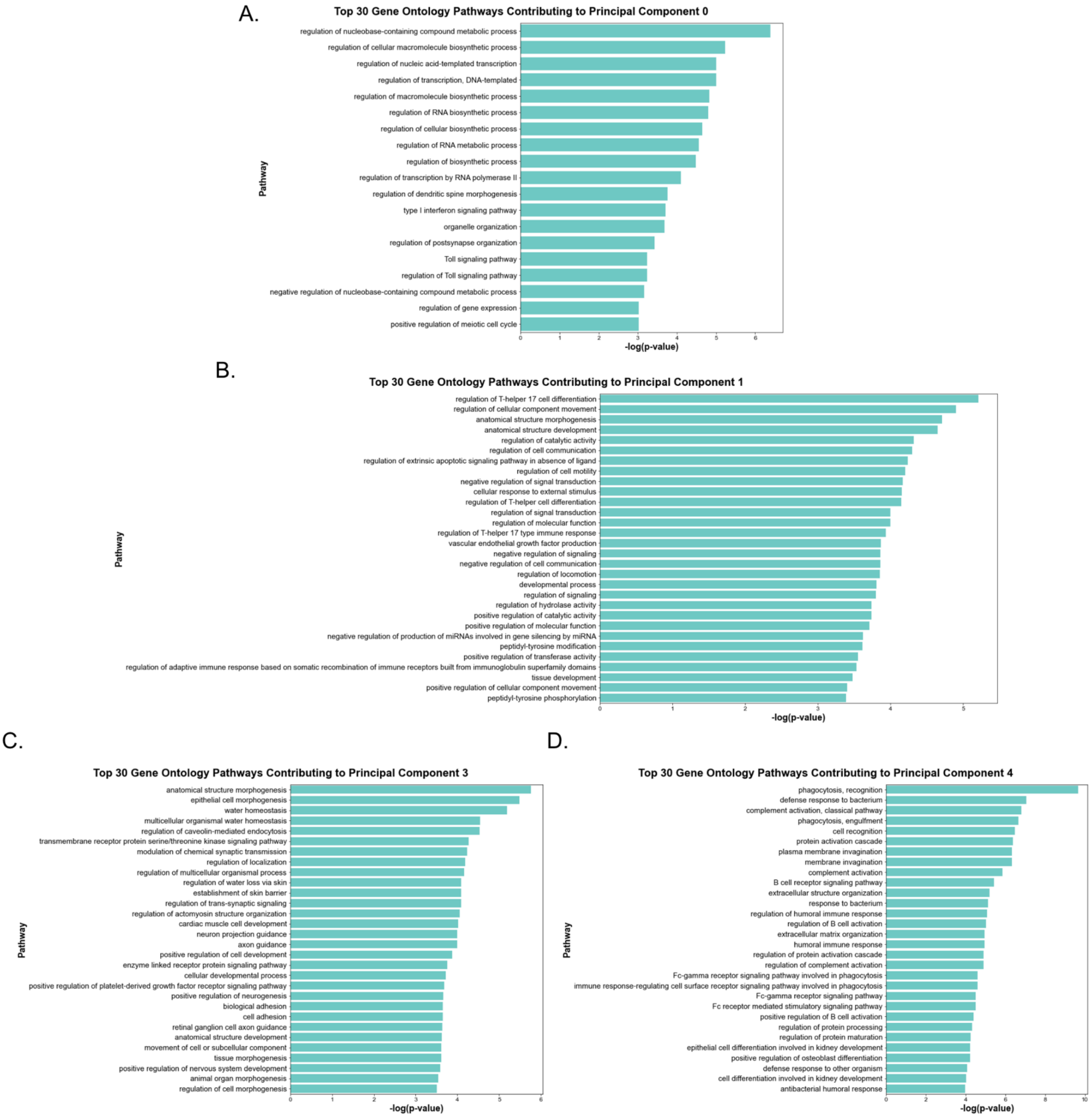


***Supplemental Figure 11. Principal Component Analysis and Associated Gene Ontology Pathways. (A-D.)*** *Bar plots of the top 30 Gene Ontology (GO) pathways for each component. PC 2 shown in main text (Figure 5). Pathways are ranked by the p-values, and significance is indicated on a logarithmic scale.*

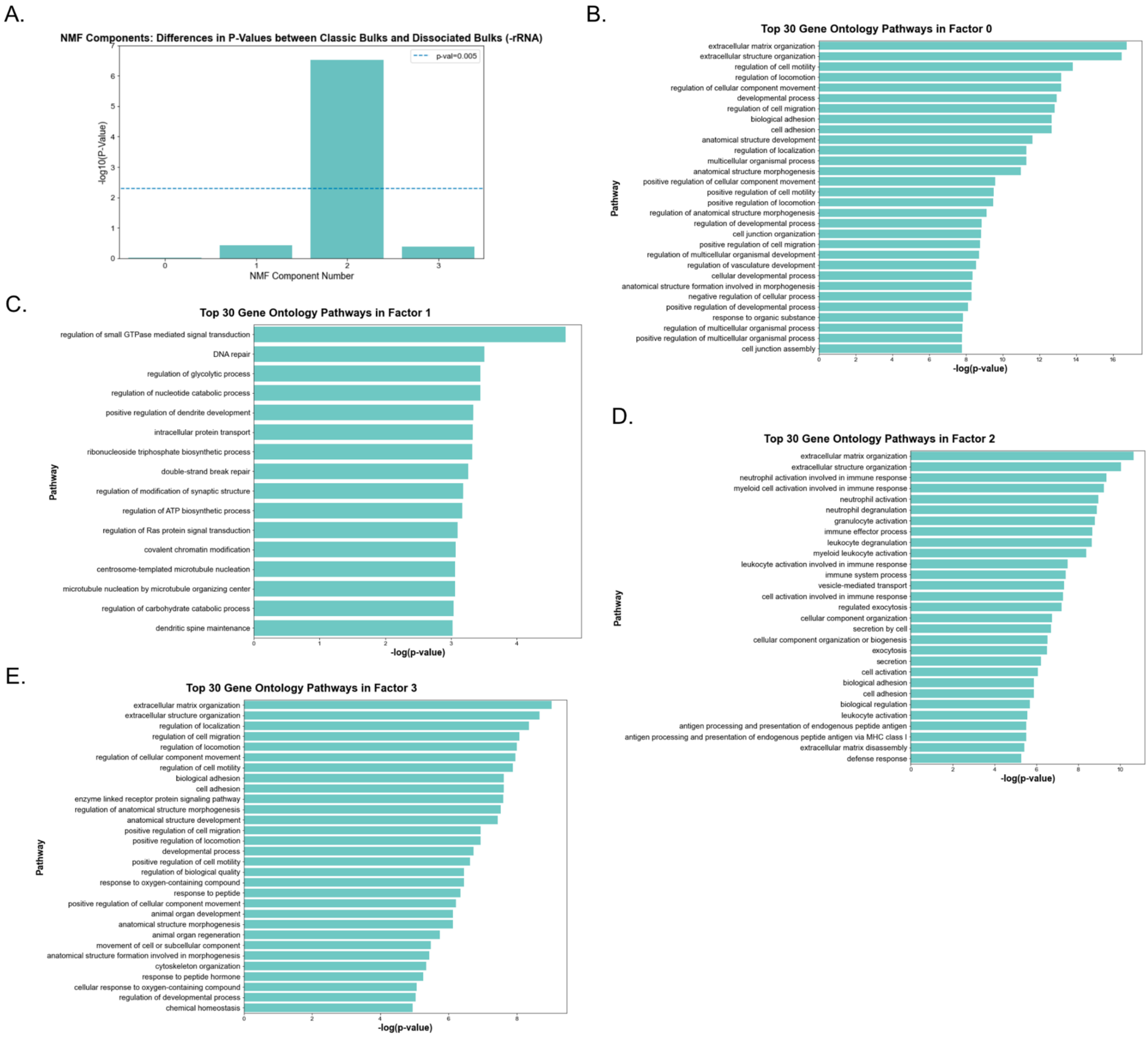


***Supplemental Figure 12. Non-negative Matrix Factorization (NMF) Analysis and Associated Gene Ontology Pathways. A.*** *Bar plot illustrating the differences in p-values between classic bulks and dissociated bulks across four NMF components, with significance denoted by -log10(p-value).* ***(B-E.)*** *Bar plots of the top 30 Gene Ontology (GO) pathways for each of the four NMF factors (0-3). Pathways are ranked by the p-values, and significance is indicated on a logarithmic scale.*
